# Circular viral copy DNA of Dengue virus (cvcDENV-2) isolated from infected mosquito cell cultures

**DOI:** 10.1101/2023.06.07.544120

**Authors:** Warachin Gangnonngiw, Wilawan Thongda, Malinee Bunnontae, Nipaporn Kanthong, T.W. Flegel

## Abstract

Dengue virus serotype 2 (DENV-2) is a mosquito-borne disease in the family *Flaviviridae*. It has been previously shown that DENV-2 can infect C6/36 mosquito cells and cause initial cytopathic effects that dissipate upon serial split-passage to yield persistently infected cultures with normal growth and morphology. In other words, the cell line accommodated persistent DENV-2 infections. It has recently been found that insect viral infections induce the production of viral copy DNA (vcDNA) fragments via host reverse transcriptase (RT). The vcDNA occurs in both linear (lvcDNA) and circular (cvcDNA) forms and produces small interfering RNA (siRNA) transcripts that can result in an immediate protective RNA interference (RNAi) response. The vcDNA can lead to the host acquiring endogenous viral elements (EVE) in genomic DNA. Thus, we hypothesized that DENV-2 cvcDNA and DENV-2-EVE would arise in C6/36 insect cells challenged with DENV-2 virus *in vitro*. Here we describe the successful isolation and characterization of cvcDNA constructs homologous to DENV-2 from laboratory challenges with C6/36 cells. At least 1 of these appeared to arise from a DENV-2-EVE. We also show that a cvcDNA preparation derived from the DENV-2 infected in C6/36 insect cells and subjected to rolling circle amplification (RCA) significantly reduced DENV-2 replication when applied to naive C6/36 cells prior to the DENV-2 challenge. This preliminary work lays the groundwork for further studies using the C6/36 cell model to screen and characterize protective EVE against insect and shrimp viruses.

## 1. INTRODUCTION

It was formerly believed that invertebrates were incapable of specific adaptive immune responses and depended solely on generic responses via innate immune mechanisms (Cerenius et al. 2010). It is well known that insects and crustaceans lack a mechanism for a specific adaptive immune response via antibodies. However, they do have a specific adaptive immune response against viruses via nucleic acids. A major feature of this response is persistent, relatively innocuous viral infections. The outcome of this response is usually persistent, but relatively innocuous viral infections. The response has been called viral accommodation and has been recently reviewed (Flegel 2020, 2022), as has similar work related specifically to insects (Bonning & Saleh 2021). To summarize, endogenous host reverse transcriptase (RT) recognizes and binds with viral RNA to produce viral copy DNA (vcDNA) in both linear (lvcDNA) and circular (cvcDNA) forms. This produces small interfering RNA (siRNA) that results in individual cell and systemic RNA interference (RNAi) responses. At the same time, some of the vcDNA fragments may be integrated into the cell genomic DNA as endogenous viral elements (EVE). The EVE may also produce antisense RNA, leading to an RNAi response. In addition, the EVE may give rise to cvcDNA, a feature that allows for its detetion in uninfected individuals because of the relative ease of isolating and sequencing it (Taengchaiyaphum et al. 2021).

It has been previously shown that C6/36 insect cells can be sequentially infected with *Aedes albopictus* densovirus (AaDNV), Dengue virus serotype 2 (DENV-2), and *Japanese encephalitis* virus to result in individual cells with innocuous, persistent triple infections that can be maintained by serial split passage (6). The mechanisms underlying the ability of these cells to accommodate 3 viruses without signs of cytopathic effects (CPE) or other signs of disease was unknown at that time. More recent work described in the previous paragraph suggested that induction of host RT and production of vcDNA might be involved. Thus, in this communication, we show that cvcDNA homologous to DENV-2 can be isolated and characterized from C6/36 cells challenged with DENV-2. We also show that DENV-2-EVE arose in the challenged, cultured cells and that these cvcDNA fragments obtained from the DENV-2 infected in C6/36 insect cells and subjected to rolling circle amplification (RCA) could significantly reduce the severity of DENV-2 infection in pre-exposed, naive C6/36 cells subsequently challenged with DENV-2.

## 2. MATERIALS AND METHODS

### 2.1 Cell cultures for DENV-2 infection

DENV-2 persistent cultures were established, as previously reported (Kanthong et al. 2008, 2010). Initially, naive C6/36 cells at 10^5^ cells/well were cultured in 6-well plates with 2 ml fresh Leibovitz’s (L15) medium containing supplements of 10% fetal bovine serum (FBS), 0.3% tryptose phosphate broth (TPB), and 1% antibiotics and incubated for 24 hours in 28°C incubator. After 24 hours of incubation, the old medium was removed from the C6/36 cell layer that was rinsed with 1 ml fresh L15 medium without other supplements. The cell layers were then challenged with DENV-2 stock at 10^5^ copies/well in 1 ml L15 medium without supplements and gentle swirling incubation at room temperature (RT= ∼25 °C) for 2 hours. After 2 hours, the 1 ml of L15 medium containing DENV-2 was removed, and 2 ml fresh L15 followed by further incubation in a 28°C incubator. Control cells were treated in the same manner, except for no exposure to DENV-2. All cell cultures were then subjected to serial sub-culture every 3 days for 7 passages (total 21 days). Each passage consisted of creating a 2 ml cell suspension divided into two 1 ml portions, one for analysis and the other for ongoing cell culture. Thus, there were 6 replicates for the test and control samples for each passage. The collected cells were stored in a -80 °C freezer until DNA extraction.

### 2.2. Circular DNA extraction and verification

Before DNA extraction, each 1 ml of suspended cells was centrifuged at 10,000 rpm for 5 min at room temperature to spin down cells. Cell samples from passages 1-7 were separately extracted for the total DNA using a DNA Extraction kit (GeneAll, Korea). The total DNA extracts of each passage were measured for DNA quantity and quality using NanoDrop™ 2000/2000c Spectrophotometers (Thermo Fisher Scientific, USA). The exact amount of 2,000 ng DNA extract from each passage (7 passages) was pooled and processed to prepare circular DNA. Total DNA was initially exposed to exonuclease V, which digests linear single-stranded-and double-stranded DNA while circular DNA remains intact. The residual putative circular DNA was isolated, purified, and collected using a DNA clean-up kit (NucleoSpin® Gel and PCR Clean-up Kit, Macherey-Nagel, Germany) following the manufacturers’ instructions.

To verify linear DNA digestion in the exonuclease V reaction step to isolate C6/36 circular DNA, we used linear DNA derived from the pGEM vector cut with TstI restriction enzyme in the same DNA amount of the C6/36 DNA and compared this to similar samples contatining ordinary pGEM vector (circular DNA). C6/36 DNA preparations containing the linear pGEM vector gave negative PCR test results for pGEM while samples containing the circular form gave positive test results. Exonuclease V enzyme was added to the reaction until linear DNA was eliminated as evidenced by no linear DNA in gel electrophoresis (1.5% agarose gel) while the circular pGEM vector was unaffected (data not shown).

Another step to confirm the successful circular DNA extraction after the exonuclease V enzyme digestion step was negative PCR test results for C6/ 36 cell genes using specific primers (**Table 1**) via polymerase chain reaction (PCR) followed by gel electrophoresis. These included use of primers for *Aedes* elongation factor gene ( EF165_*Aedes*_F/ R, Table 1) . In contrast, positive PCR test results were obtained using mitochondrial ( mt) DNA genes of the *Aedes* ATP Synthase and cytochrome c oxidase primers ( ATPSynthase_150F/ R and CYCS_*Aedes*_125F/R, Table 1).

**Table 1.**
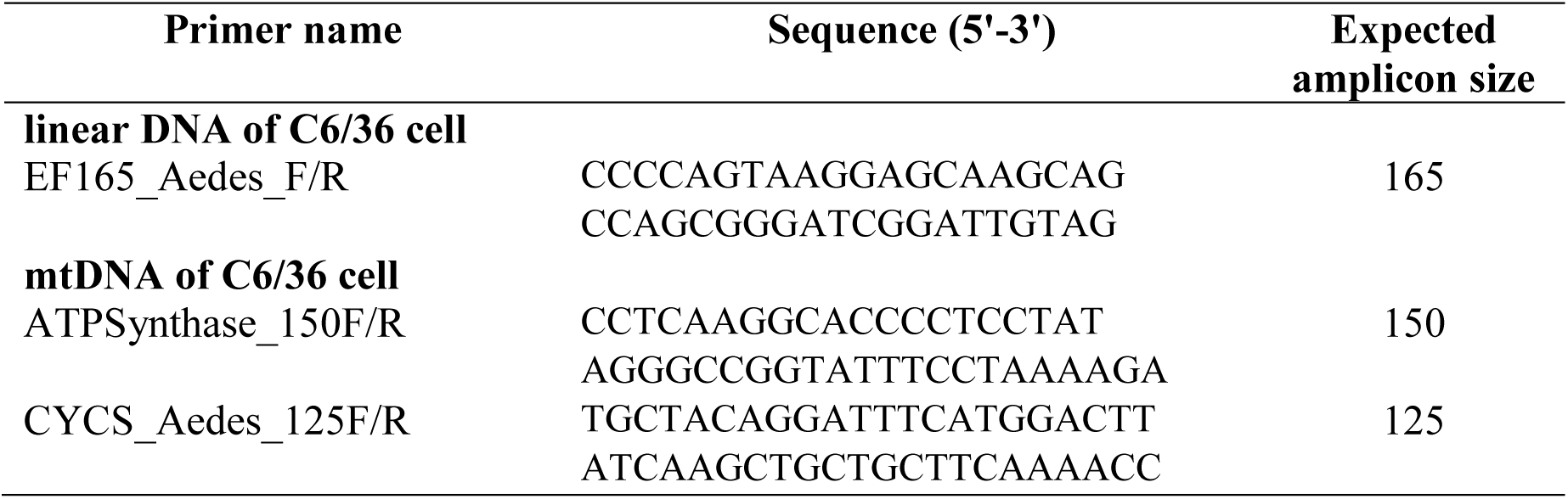
Primers were used to amplify genes of C6/36 cells in this study.

### 2.3. Sequencing of cvcDNA

To verify the result of cvcDNA of DENV-2 identification, the pooled circular DNA extracts after the exonuclease V enzyme digestion were subjected to metagenomic sequencing. The first pooled circular DNA isolation was submitted directly to GENEWIZ, Inc., Suzhou, China, for library construction and sequencing without any PCR amplification step. A second pooled circular DNA preparation was subjected to circular DNA amplification using Phi29 DNA polymerase rolling circle amplification (RCA) (TempliPhi™ 100 Amplification Kit, Cytiva, USA) according to the manufacturer’s instructions. This second pooled circular DNA sample was submitted to NovogeneAIT Genomics, Singapore, for library preparation and metagenomic sequencing. The specimens submitted were in the amount (Qubit®) ≥ 200 ng, volume ≥ 20 µl, concentration ≥ 10 ng/ µl, and purity (Nanodrop^TM^/Agarose Gel) OD260/280 = 1.8-2.0. Both companies confirmed that there was no degradation or contamination and that the samples passed quality and quantity requirements for DNA sequencing.

### 2.2 Identification of cvcDNA of DENV-2 from circular DNA sequencing

Circular DNA library preparation and sequencing were separately performed by GENEWIZ, Inc., Suzhou, China (1^st^ pooled DNA extracts without any PCR amplification step), and NovogeneAIT Genomics, Singapore (2^nd^ pooled DNA extracts after rolling circle amplification (RCA)). A TruSeq DNA Library Preparation Kit (Illumina) was used for cDNA library preparation, following the TruSeq protocol. Briefly, DNA was fragmented by sonic disruption using a Covaris S220 apparatus (Covaris). Subsequently, blunt-end fragments were created, and the A-base was tailed to each 3’side. Dual indexes within P5 and P7 adapters were ligated to the fragments. The final products were purified. Then, the size of the purified DNA was selected before PCR amplification for library construction. Initial DNA concentration was quantified using Qubit 3.0, and fragment size was determined using Agilent 2100. The final library samples were additionally examined by Agilent 2100. Afterward, amplified libraries were sequenced on an Illumina HiSeq 2000 instrument with a 150 bp pair-end (PE).

The raw reads were generated in FASTQ format. The company, moreover, cleaned up raw reads by removing the adapter sequences. The reads with low-quality scores and those containing ambiguous nucleotides (N) greater than 10 percent were removed. Clean reads were obtained for further bioinformatics analysis. The clean raw reads were *de novo* assembled into contigs using MEGAHIT (Version 1.2.9) by Li et al. (2015) under the Galaxy platform (The Galaxy Community 2022). Subsequently, the assembled contigs were used as query sequences to search BLASTN against the dengue virus serotype 2 (DENV-2) complete genome database (accession no. M29095.1; 10723 bp ss-RNA). BLASTN (NCBI BLAST+ blastn). The program was run under the Galaxy platform (The Galaxy Community 2022) with a cutoff E-value of 1x10^-30^.

The hits between contigs and DENV-2 (M29095.1) from BLASTN output were retrieved for sequences that were further analyzed for the presence of cvcDNA. The retrieved sequences were performed using the program Filter FASTA on the sequence headers with a list of IDs. Afterward, the sequences that matched with DENV-2 (M29095.1) were further analyzed for *Aedes albopictus* isolate C6/36 fragments. To detect *Aedes albopictus* isolate C6/36 fragments in the retrieved contig sequences, the BLASTN program was applied using *Aedes albopictus* isolate C6/36 whole genome (GCF_001876365.2) as the database (subject sequences). Multiple sequence alignments were performed using CLUSTALW (https://www.genome.jp/tools-bin/clustalw). The circular DNA diagrams were drawn by GenomeVx (http://wolfe.ucd.ie/GenomeVx/). Moreover, the clean raw reads by GENEWIZ, Inc. were analyzed for similarity to DENV-2 sequences using map reads and the reference program QIAGEN CLC Genomics Workbench version 12 (QIAGEN, Denmark) to reconfirm the similarity of raw reads and contig assembly compared to DENV-2 genome.

### 2.3 Validation for the presence of DENV-2 contigs in the circular DNA preparation

Multiple specific single primers were designed from selected contig sequences to confirm DENV-2 cvcDNA in the persistently infected DENV-2 cultures (or circular DNA extract). PCR amplification with each primer using a portion of the original, pooled circular DNA sample as a template (30 ng) was carried out using KOD FX Neo polymerase (Toyobo, Japan), following the manufacturer’s instructions for PCR conditions. The thermal cycling profile consisted of an initial denaturation at 98° C for 2 minutes, followed by 35 cycles of denaturation at 98° C for 10 seconds, appropriate annealing temperature at 60 °C for 5 seconds, and extension at 68°C for 1 minute. The PCR products were visualized by 1.5% agarose gel electrophoresis. There was no confirmation by submitting PCR products for sequencing again.

### 2.4 Anti-DENV-2 activity tests using circular vcDNA (cvcDNA) produced by RCA

For the anti-DENV-2 activity test (**Figure 1**), naive C6/36 cells at 10^5^ cells/well were seeded into 6-well plates with 2 ml fresh Leibovitz’s (L15) medium with supplements containing 10% FBS, 0.3% TPB, and 1% antibiotics and incubated for 24 hours at 28°C before transfection using liposome Escort IV transfection reagent (Sigma) according to the manufacturer’s instructions. After 24 hours of incubation, the medium was removed and the cells were rinsed with 1 ml fresh L15 medium without supplements. Then, 800 µl fresh L15 cell culture medium without supplements was added. At the same time, the transfection reagent was separately prepared by (A) 4 μl Escort IV transfection reagent mixed in 100 μl L15 medium without supplements and (B) 100 or 1000 ng of cvcDNA mixed in 100 μl L15 medium without supplements and then mixing (A) and (B) to form liposome-forming complexes for 45 minutes at room temperature. Then, the ∼200 μl mixtures (A and B) were added dropwise to the cell monolayers in 800 μl fresh L15 medium without supplements with gentle swirling for 5 hours at room temperature. After 5 hours of incubation, they were challenged with DENV-2 stock at 16.85 x 10^6^ copies/well in 1 ml of L15 medium without supplements followed by gentle swirling for 2 hours at room temperature. After incubation for 2 hours, the old medium was removed, and 2 ml fresh L15 medium with supplements (10% FBS, 0.3% TPB, and 1% antibiotics) was added for further incubation at 28°C until day 5.

**Figure 1.**
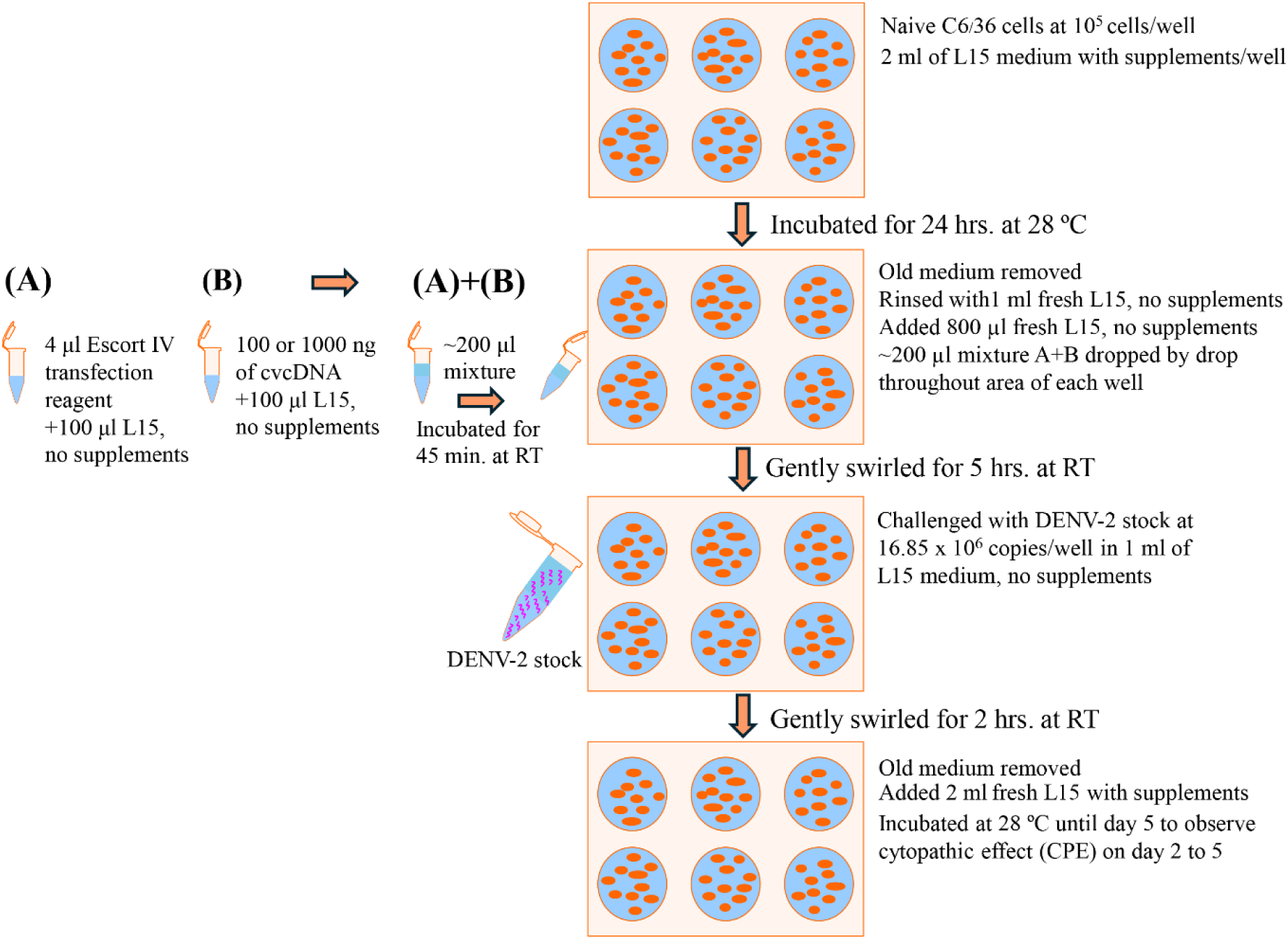
Diagram of the protocol for anti-DENV-2 activity tests using circular vcDNA.

An experiment to test for anti-DENV-2 activity was carried out with 4 groups, including 1) negative control (naive C6/36 cells), 2) positive control (C6/36 cells+DENV-2), 3) treatment of 100 ng cvcDNA (C6/36 cells+100 ng cvcDNA+DENV-2), and 4) treatment of 1,000 ng cvcDNA (C6/36 cells+1,000 ng cvcDNA+DENV-2) with six replicates in each group. The presence of any cytopathic effect (CPE) was assessed on days 3 to 5 to examine the severity of viral pathology. Pictures of CPE were captured using an inverted microscope (Leica DMi1) with 20X magnification. Three pictures of each replicate were captured. The seriousness of CPE was assessed using a grid of 100 squares in one picture. The infected area was counted as 1 for squares containing syncytial cells and as 0 for squares without CPE. The total infected areas were then summed. Statistical analysis was performed using the R program, with a one-way ANOVA followed by the Tukey HSD post hoc test (***p-value ≤ 0.001, ns = non-significant).

## 3. RESULTS

### 3.1 Confirmation of circular DNA isolation from DENV-2 infected C6/36 cell cultures

To verify the extraction of circular DNA from DENV-2 infected C6/36 cell cultures by our isolation process, DNA extracts before and after exonuclease V digestion were examined for the presence of linear DNA and circular DNA using PCR amplification with specific primers followed by visualization with gel electrophoresis. Linear DNA was detected using the primers EF165_Aedes_F/R (Table 1) that amplified host elongation factor gene. Circular DNAs were detected using two primers of ATPSynthase_150F/R and CYCS_Aedes_125F/R (Table 1) that amplified mtDNA genes of ATP synthase and cytochrome c oxidase, respectively. The result in **Fig. 2**. showed the presence of linear DNA amplicons in lane 1 (positive amplification confirming presence) derived from elongation factor (EF) gene primers before DNA extracts were treated with exonuclease V. These amplicons were absent after DNA extracts digested with exonuclease V (lane 4 negative amplification confirming absence). In contrast, there were amplicons present for the circular DNAs from mtDNA genes (ATP synthase and cytochrome c oxidase) both before (lanes 2 and 3) and after (lanes 5 and 6) exonuclease V digestion (Figure 2). These results confirmed that only host linear DNA was eliminated while circular DNA forms remained. The presence of circular DNA initially supported the proposal for presence of a cvcDNA construct corresponding to the dengue virus single-stranded positive-sense RNA genome. Therefore, metagenomic sequencing was subsequently conducted to reveal the sequence in the circular DNA in the cvcDNA preparation.

**Figure 2.**
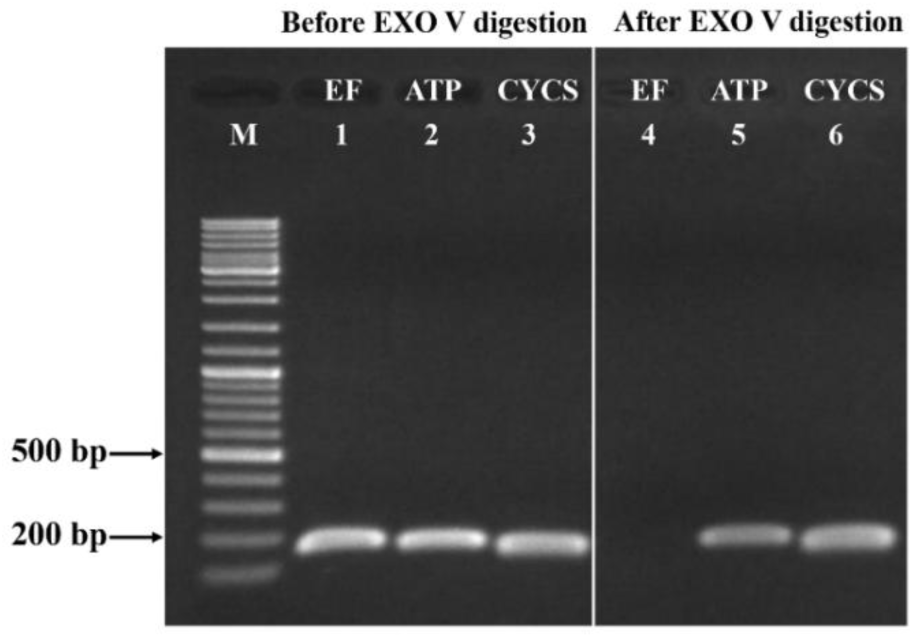
Analysis of PCR assays by agarose gel electrophoresis to confirm host circular DNA presence but linear DNA absence following exonuclease V (EXO V) digestion. Before EXO V digestion, Lane 1 shows amplicons for the linear DNA of host elongation factor gene (EF, Lane 1) and for two mitochondrial circular DNA genes, ATP synthase (ATP Lane 2) and cytochrome c oxidase (CYCS Lane 3). After EXO V digestion, the EF amplicon is absent (Lane 4) while the two mitochondrial gene amplicons (ATP Lane 5 & CYCS Lane 6) still remain. M is a GeneRuler DNA ladder mix.

### 3.2 Identification of DENV-2 cvcDNA from circular DNA metagenomic sequencing

#### 3.2.1 Raw reads matching DENV-2 mapped irregularly across its whole genome

Based on the viral accommodation concept, we hypothesized that the RNAi and host reverse transcriptase activities would generate vcDNA with high variation in size and sequence and that a portion of this would be in circular form (i.e., cvcDNA), even though DENV-2 is an RNA virus. Metagenomic sequencing was utilized to test this hypothesis. To ensure the next-generation sequencing procedure, we submitted two separate circular DNA extracts to two companies (see section Materials and Methods 2.2). The raw reads of two metagenomic sequencings were deposited as NCBI short read archive (SRA) under BioProject of PRJNA1003297 and PRJNA1160223 that are raw reads from GENEWIZ, Inc. with the ID of D2C6P1-7, and raw reads from NovogeneAIT Genomics with the ID of CCDENV21. The sample of D2C6P1-7 generated 29,182,977 pair-end (150 bp) raw reads, while the sample of CCDENV21 generated 35,174,410 pair-end (150 bp) raw reads.

From each company, raw reads that matched the DEN-2 genome sequence were mapped to the reference genome M29095 of DENV-2 (**Fig. 3**). Distribution of the DENV-2-matching raw reads was irregular and spanned across most of the genome sequence. This is consistent with the proposal that EVE acquisition is an individual cell process that leads to high variation in the types and numbers obtained from a tissue sample or cell culture. In contrast, raw data from circular DNA preparation (D2C6P1-7) revealed a high-frequency distribution of genes only within the ENV-NS2A region of the DENV-2 genome. This roughly matched a similar map region for the CCDENV21 data set.

**Figure 3.**
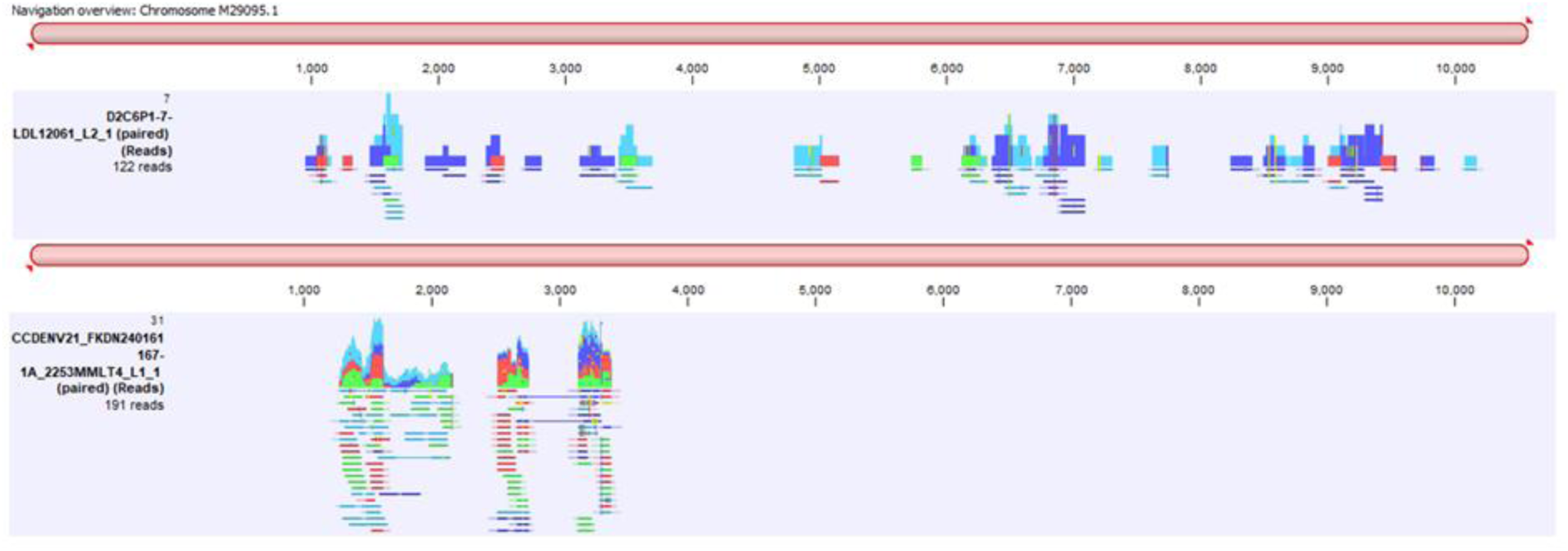
Diagram showing the number, length and distribution of 122 raw reads obtained from the metagenomics of the D2C6P1-7 (122 raw reads) and CCDENV21 (191 Raw reads) cvcDNA samples when mapped against the reference DENV-2 genome (GenBank M29095).

Both raw read samples were separately assembled using MEGAHIT, yielding 110,350 and 450,310 contigs for D2C6P1-7 and CCDENV21, respectively. Given the emphasis on DENV-2 cvcDNA, we further analyzed the contigs from two samples using BLASTN. Each contig assembly was tested separately using BLASTN against the dengue virus serotype 2 (DENV-2) complete genome database (M29095.1). With a cut-off E-value of 1x10^-10^ when using the BLASTN program, the results showed that the D2C6P1-7 sample with 110,350 contigs yielded 3 contigs that matched the DENV-2 sequence. These were k141_3180 (368 bp), k141_92607 (433 bp), and k141_113249 (327 bp) (supplementary data **Table S1**).

BLASTN output of the CCDENV21 sample with 450,310 contigs showed 5 DENV-2 matching contigs that included contigs k141_157363 (417 bp), k141_311629 (281 bp), k141_42362 (298 bp), k141_43783 (2061 bp), and k141_554965 (499 bp) (supplementary data **Table S1**). Sequences of all eight contigs were retrieved for further comprehensive analysis (supplementary data **Table S2**).

The 8 contig sequences were subsequently analyzed for host DNA fragments (*Aedes albopictus* isolate C6/36) using BLASTN. This process was run to confirm the presence of DENV-2 and *A. albopictus* isolate C6/36 sequences together in the same contig. The BLASTN output showed three contigs containing sequences of dengue virus serotype 2, joined with fragments of DNA from the *A. albopictus* isolate C6/36 (supplementary data **Table S3**). All three contig sequences, including contigs k141_157363, k141_42362, and k141_43783, were derived sequences, including contigs k141_157363, k141_42362, and k141_43783, were derived from the CCDENV21 sample. The feature summary of raw reads, contigs from raw reads assembly, the retrieved contig sequences containing the DENV-2 sequences, and the *Aedes albopictus* isolate C6/36 fragments from each company are shown in **Table 2**.

**Table 2.**
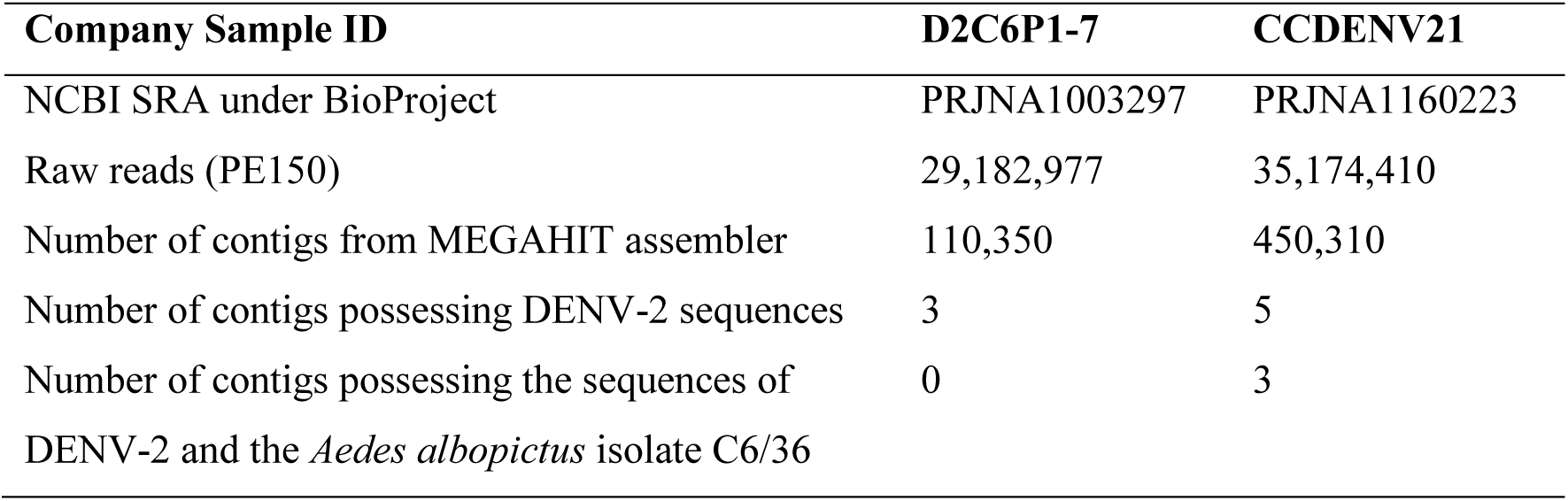
The feature summary of raw reads and the number of contigs from the raw read Assembly.

#### 3.2.2 Characterization of eight contigs of DENV-2 cvcDNA

As mentioned above, cvcDNA constructs would vary in size and sequence. BLASTN results of **Table S3** demonstrated that the sequences of *A. albopictus* isolate C6/36 in the cvcDNA contigs may have originated from several locations in the host genome. For example, the sequences at 278-417 (140 bp) of contig k141_157363 and the sequence at positions 1-140 (140 bp) of contig k141_42362 gave hits at 49 and 47 different locations in the *A. albopictus* isolate C6/36 genome (supplementary data **Table S3**). Contig k141_43783 showed positions at 387-1026 (640 bp), 387-1031 (645 bp), 388-603 (216 bp), 1028-1122 (95 bp), 1029-1117 (89 bp), and 1110-1194 (85 bp) that mapped to 19 locations in host genome (supplementary data **Table S3**). These results indicated that DENV-2 cvcDNA contained a wide variety of viral and host gene fragments in variable numbers.

Subsequently, all eight contig sequences were examined for circular forms by searching for overlaps within each sequence. The results showed that contigs k141_43783 (2061 bp) and k141_554965 (499 bp) had complementary overlap sequences of 141 bp at the ends of their contigs (supplementary data **Document S1, Figure S1**). In **Table 3**, in the far-right column, the reading directions within the contigs are indicated by arrows with reference to the DENV-2 5’V3’ genome sequence, and the vertical bars indicate lack of continuity in the sequences and reading directions. These features, especially when accompanied by linked host DNA sequences, support the contention that the contigs arose from cvcDNA constructs derived from the host genome and not directly from the DENV-2 genome. The circular diagram of two contigs of DENV-2 cvcDNA is illustrated in **Fig. 5**. The 141 bp overlap regions were merged. Thus, the contigs k141_43783 and k141_554965 represent cvcDNA constructs of 1921 and 358 bp, respectively.

**Table 3.** Summary features for eight contigs that hit the DENV-2 and/or *Aedes albopictus* 346 C6/36 cell genome sequences. The contigs derived from raw read set D2C6P1-7 are highlighted 347 in a gray background, while those derived from set CCDENV21 are not highlighted. In the 348 column for “Position and Reading Direction”, arrows indicate reading direction and vertical 349 bars indicate disjunction between the DENV-2 sequence connections. Yellow highlights in the column entitled “Fragment Positions” indicate overlaps (i.e., common bases at the junction of two fragments).

| # | Contig Name | Length (bp) | C6/36 content | Circular (yes/no) | # Fragments | Fragment Positions | DENV-2 Genome Position and Reading Direction (← →) |
| --- | --- | --- | --- | --- | --- | --- | --- |
| 1 | k141_3180 | 368 | No | No | 3 | 26-253 (228)<br>248-323 (76)<br>321-368 (48) | 1458-1685→<br>\| 1246-1321→ \|<br>1082-1129→ |
| 2 | k141_92607 | 433 | No | No | 2 | 13-128 (116)<br>116-413 (298) | ← 7302-7188 \|<br>6374-6671→ |
| 3 | K141_113249 | 327 | No | No | 1 | 21-327 (307) | 1287-1567→ |
| 4 | K141_157363 | 417 | Yes | No | 3 | 1-147 (147)<br>147-281 (135)<br>278-417(140) | ← 1422-1276 \|<br>← 1619-1485<br>C6/36 seq |
| 5 | K141_311629 | 281 | No | No | 1 | 1-281 (281) | 1287-1567→ |
| 6 | K141_42362 | 298 | Yes | No | 2 | 1-140 (140)<br>141-158 (18)<br>159-298 (140) | C6/36 seq<br>No identity<br>1485-1346→ |
| 7 | K141_43783 | 2061 | Yes | Yes | 6 | 1-141 (141)<br>141-387 (247)<br>387-1031 (645)<br>1028-1122 (95)<br>1110-1194 (85)<br>1179-2061 (883) | ← 1422-1282 \|<br>← 2758-2512 \|<br>C6/36 seq.<br>C6/36 seq.<br>C6/36 seq.<br>← 2165-1282 |
| 8 | K141_554965 | 499 | No | Yes | 5 | 4-96 (93)<br>95-272 (178)<br>277-361 (85)<br>362-454 (93)<br>453-499 (47) | ← 2757-2665 \|<br>← 3319-3142 \|<br>\| 3316-3400→<br>← 2757-2665 \|<br>← 3319-3273 |

The feature summary of eight contigs in Table 3 and the BLASTN results in the supplementary data (Table S1) revealed that the eight contig sequences originated from positions scattered across the DENV-2 genome. Some contigs from two different company pools of raw reads also shared common areas. The eight contigs generated 18 fragments that, in a scrambled and scattered manner, matched their positions and reading directions in the DENV-2 genome (Table 3 and Fig. 4).

**Figure 4.**
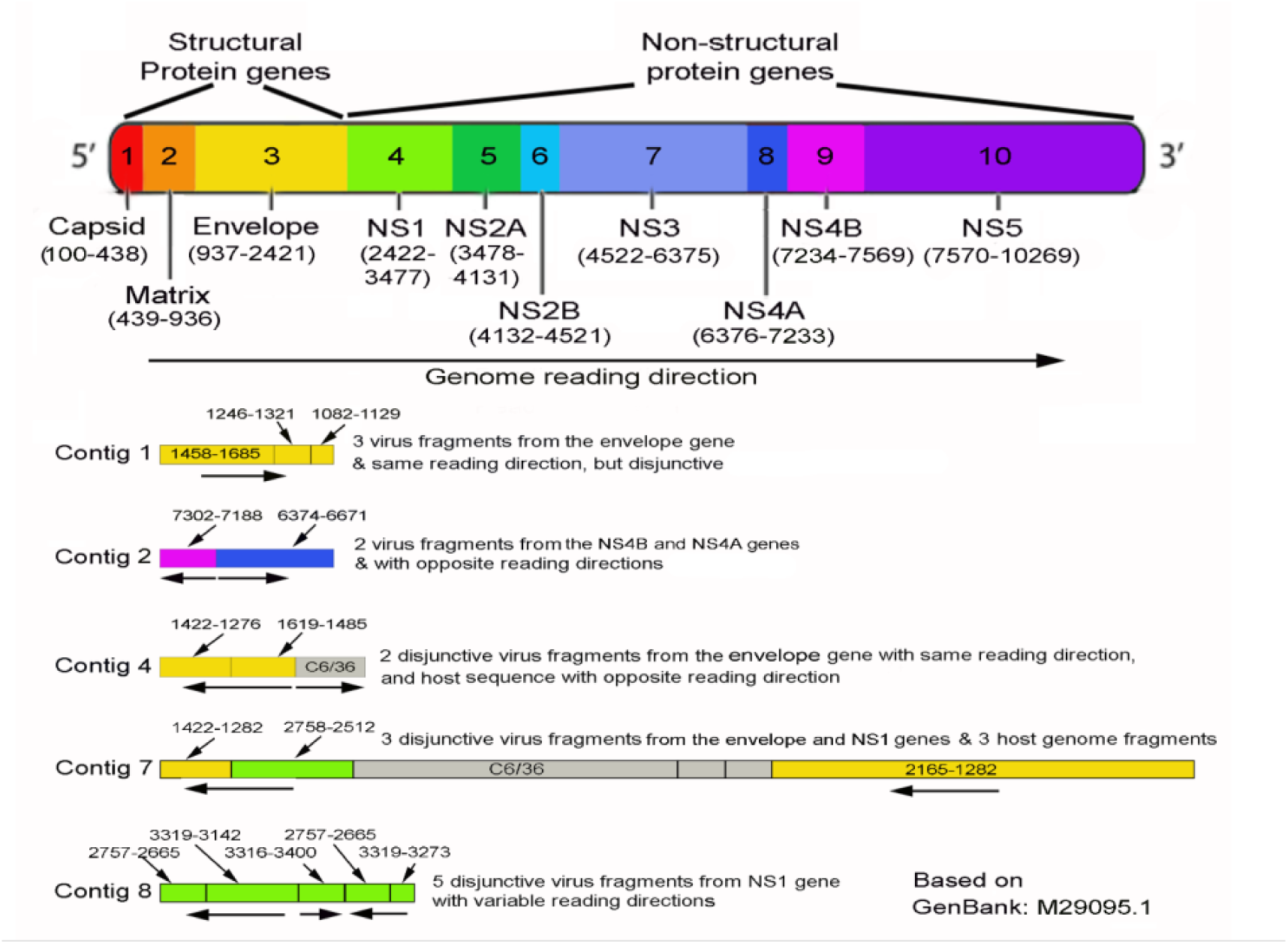
Diagram of the reference DENV-2 polyprotein genome together with five representative contigs from **Table 3**. The scales are relative, with the contigs magnified about 5 times compared to the whole genome to make the fragment features more visible. These contigs illustrate the features of chimeric gene fragment compositions (including viral genome fragments or viral plus host genome fragments), the linkage of disjunctive viral and host genome fragments, and variable reading directions (arrows below the contig bars), which can be compared to the reference genome diagram at the head of the illustration.

### 3.3 Confirmation of the presence of circular DENV-2 DNA related to contig sequences

According to **Table 3** and **Figs. 4** and **5**, the data showed that the DENV-2 genome content in the contigs originated from the region of the genome where most raw-read matches occurred at positions 1000-2000. Multiple single primers were designed to detect circular DENV-2 DNA using DENV-2 contig sequences at positions 1000-2000. The single primer of cvcDENV2_1422 5’-GATTTCCTTGCCATGTTTTCCTGTGTCA-3’ was directly designed from the sequence of contig k141_43783, and another single primer of cvcDENV2_1422x 5’-TGACACAGGAAAACATGGCAAGGAAATC-3’ was designed from the antiparallel complementary sequence of contig k141_43783. A portion of the final circular DNA preparation obtained from using RCA (the same sample submitted for Novogene sequencing). was used as a template to test these single primers. When each of these primers were used alone for PCR amplification, they successfully produced PCR amplicons (**Fig. 6**.). For example, the single primer for cvcDENV2_1422x gave several amplicon bands of different sizes, indicating the presence of a variety of circular DENV-2 DNA targets, with the most robust band found at an amplicon size of ∼500 bp in both primers and the product size of ∼2000 bp from a primer of cvcDENV2_1422 (**Fig. 6**). These results also supported the idea that cvcDNA produced from the same DENV2 genome regions were produced in variable sizes.

**Fig. 5.**
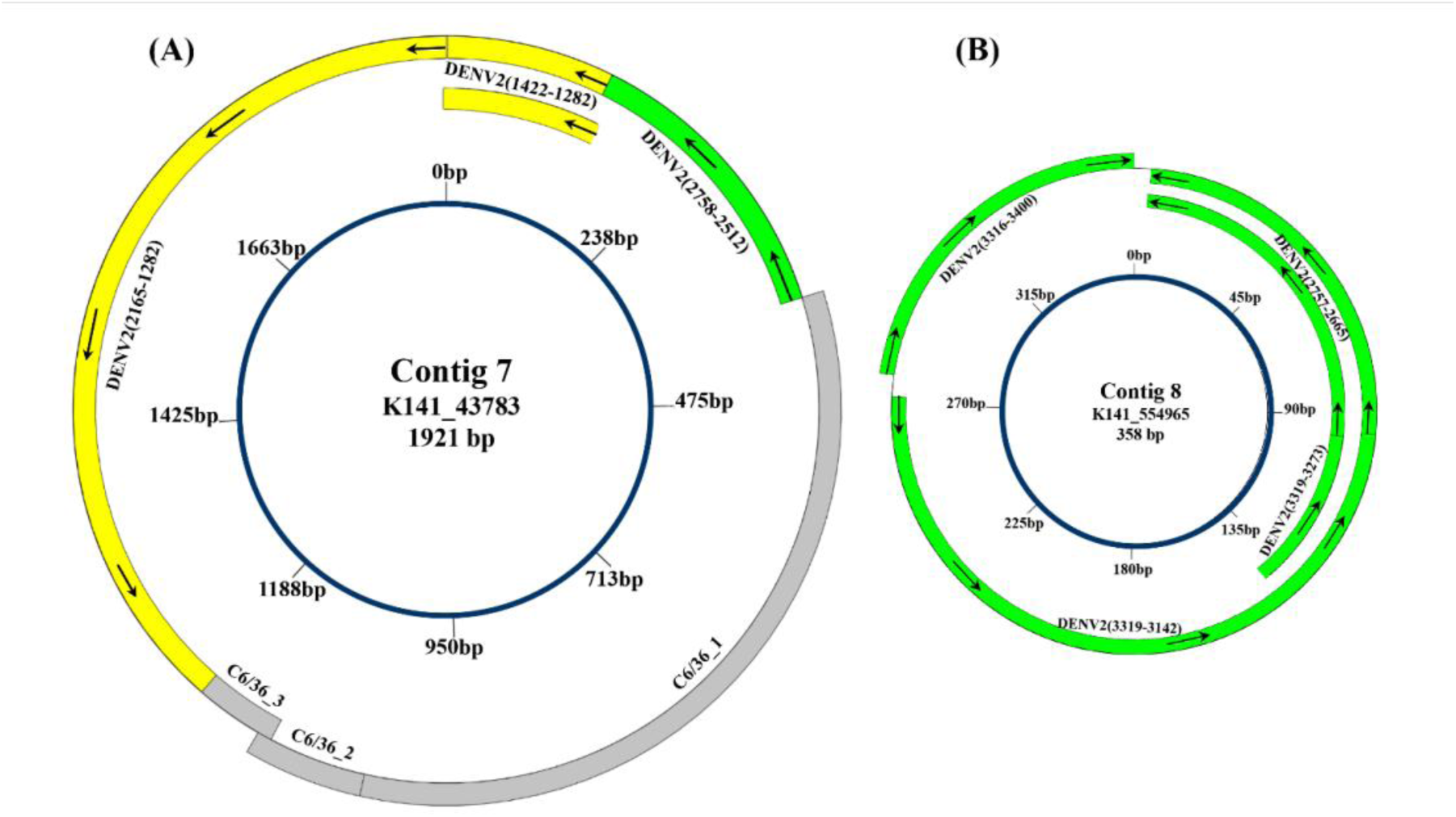
Diagram of 2 circular DNA contigs, k141_43783 and k141_554965. The grey fill 377 indicates C6/36 host DNA sequences, while the DENV-2 sequences are filled with colors to match the DENV-2 genes illustrated in **Fig. 4** (envelope gene in yellow and NS1 gene in green). **(A)** Contig k141_43783 (2,061 bp) contains three disrupted fragments of the DENV-2 envelope protein gene plus three disruptive fragments of C6/36 DNA. **(B)** Contig k141_554965 (499 bp) contains five DNA fragments derived from the DENV-2 genome (M29095), positions varying from 2,665 to 3,400. The inner blue circle indicates the position of the nucleotides in the contigs. Note the opposite reading directions in the two single contigs, each containing disruptive sequences. The inner double-circle part represents overlapping sequences, each 141 bp, resulting in contig 7 (k141_43783) with 1,921 bp and contig 8 (k141_554965) with 358 bp in circular form. The figure does not show the actual contig size scales in the comparisons, but the figure still shows that contig 8 (k141_554965) is smaller than contig 7 (k141_43783), although it is actually 5 times smaller.

**Fig. 6.**
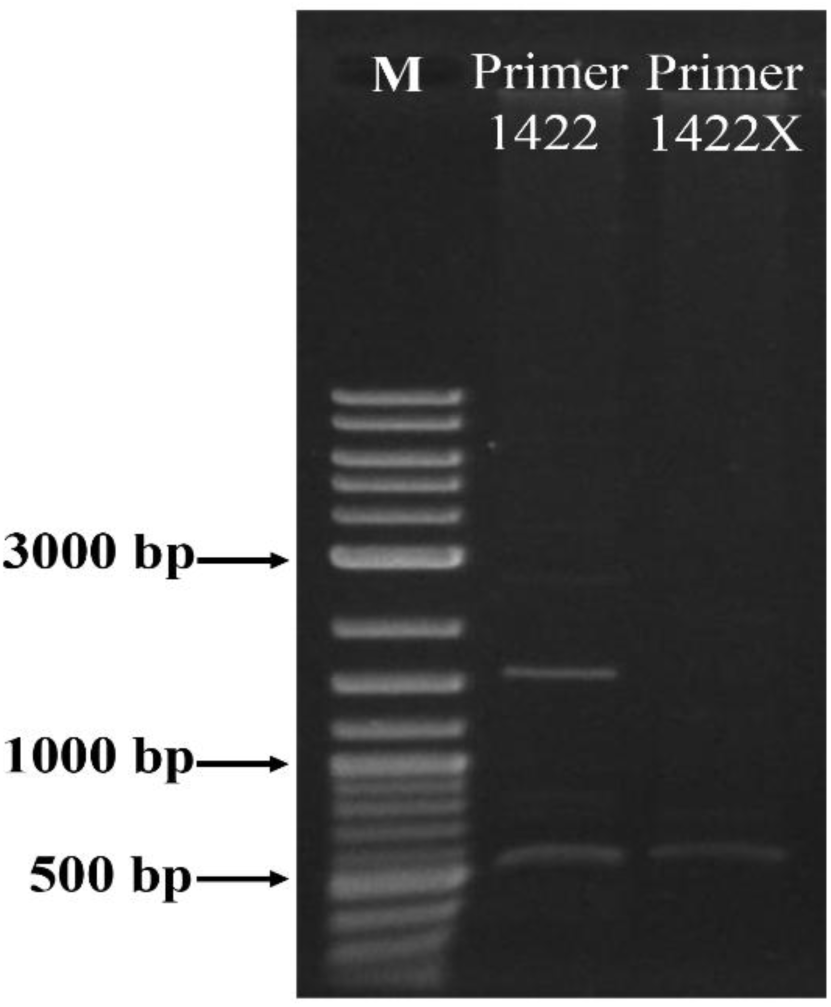
Photograph of an agarose gel showing that PCR amplicons were obtained using 2 different single primers (cvcDENV2_1422 and cvcDENV2_1422X), indicating the presence of palindromic repeats. Lane M is a GeneRuler DNA ladder mix.

### 3.4 DENV-2 tolerance in C6/36 cells accelerated by cvcDNA of DENV-2 treatment

In viral accommodation, vcDNA arises as linear (lvcDNA) and circular (cvcDNA) forms via endogenous host RT activity, leading to a viral-tolerant, persistent infection state. Here, a mixture of cvcDNA of DENV-2 obtained using the cvcDNA preparation amplified by RCA (see section 2.1 of the Materials and Methods) was used in pre-exposure tests with naive C6/36 cells prior to the DENV-2 challenge (**Fig. 1)**. The consistent outcome for naive cells challenged with DENV-2 is always a significant occurrence of CPE exemplified by syncytial cells on day 2, followed by a progressive decrease in CPE until day 5. This is shown in **Fig. 7**, where CPE can be readily observed on day 2 in untreated cells. This contrasts with the cells treated with DENV-2-cvcDNA (at 100 and 1,000 ng) on day 2, in which only a minor CPE was observed. On day 5, the untreated groups showed distinguishable morphological differences, with clearly visible, more-severe CPE in the untreated cells (**Fig. 7**) that showed. a significant reduction in CPE. This result was confirmed statistically by quantifying the infected areas, as shown in **Fig. 8**. It revealed that the presence of DENV-2-cvcDNA prior DENV-2 challenge resulted in the cells accommodating DENV-2 faster than untreated cells and confirmed it association with accommodation.

**Fig. 7.**
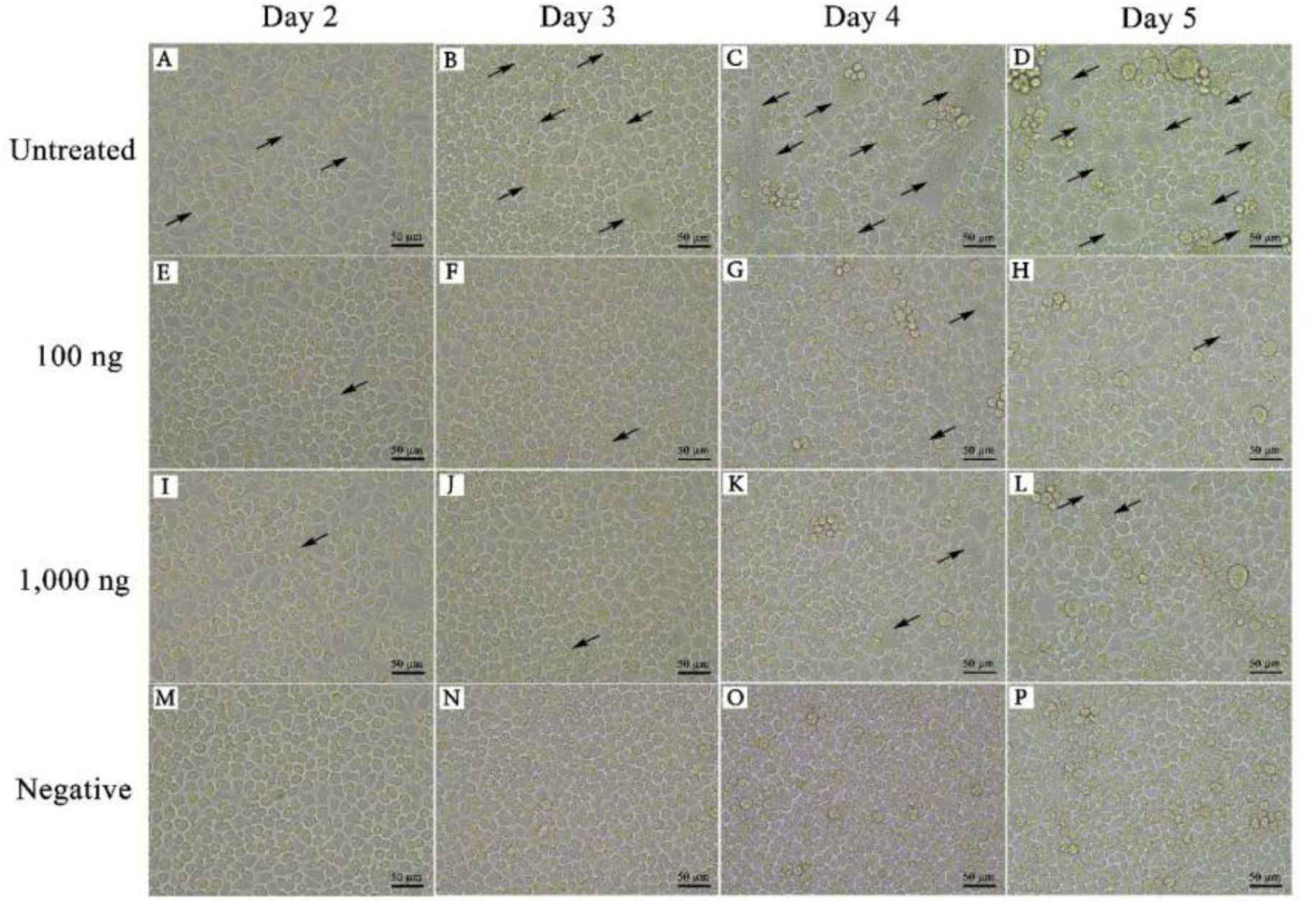
The cell morphology of DENV-2-infected cells, 2,3,4, and 5 days after infection, treated or not treated with DENV-2 cvcDNA (100 and 1,000 ng) prior to the DENV-2 challenge. CPE was observed by syncytial cells (black arrows). The untreated cells exhibit significant CPE, as evidenced by the formation of syncytia (black arrows).

**Fig. 8.**
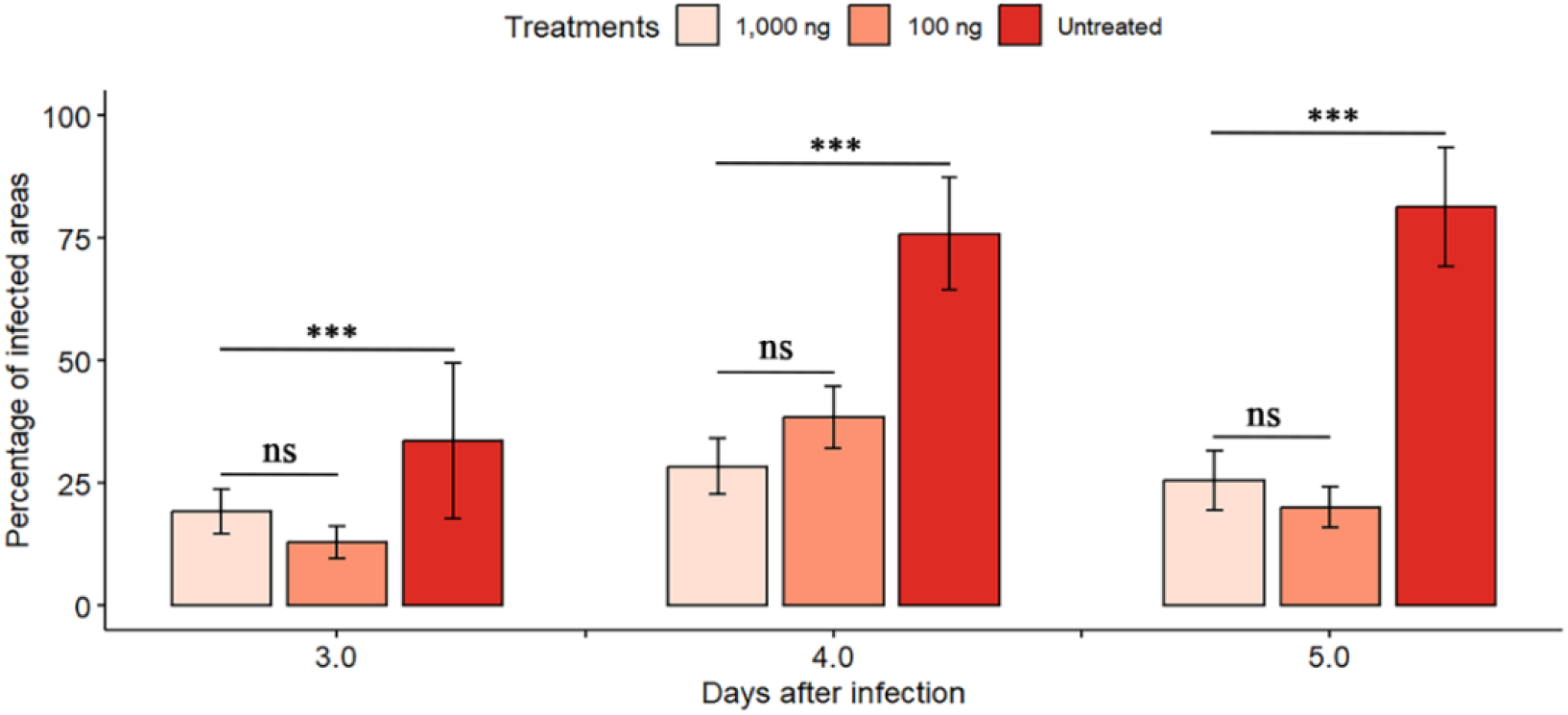
The percentage of the infected areas indicated by CPE on days 3, 4, and 5, in cells treated or not treated with DENV-2 cvcDNA (100 and 1,000 ng) prior to DENV-2 challenge. CPE was assessed by the presence of syncytial cells. Statistical analysis was performed using 447 the R program and analyzed with one-way ANOVA, followed by the Tukey HSD post hoc test (***p-value ≤ 0.001; ns = non-significant).

## 4. DISCUSSION

The tiny proportion of DNA raw reads homologous to DENV-2 (122 raw reads) in the total of over 58 million raw reads from GENEWIZ, Inc., and the 284 raw reads in 70 million raw reads from Novogene Co., Ltd, was not unexpected. However, these results were consistent with the number of contigs from the MEGAHIT assembler that showed three contigs homologous to DENV-2 obtained from a total of 110,350 contigs from GENEWIZ, Inc., and the five contigs homologous to DENV-2 derived from a total of 450,310 contigs obtained by Novogene Co., Ltd (Table 2). From **Figs. 5** and **6**, with the naked eye, there are clusters of slightly higher frequency raw reads around the region of ENV (DENV-2 positions of 937-2421), but no statistical analysis was carried out to determine whether these differences were significant. However, in this brief communication, the results supported our hypothesis that C6/36 cells infected with DENV-2 would produce cvcDNA constructs containing DENV-2 homologous sequence fragments.

As seen in **Figs. 3** and **4**, **Table 3**, and supplementary data **Table S2**, the sequences of these eight contigs could not have arisen directly from a DENV-2 viral genome because of the disjunctive and opposing reading directions of different portions of the viral genome that were connected in the contigs. In addition, three contigs contained sequences matching disjunctive fragments of the host genome sequence linked to the DENV-2 sequences. These features support the contention that circular DNA entities may have arisen from the EVE of DENV-2and been acquired after DENV-2 infection.

We have confirmed that DENV-2 infections in C6/36 cell cultures, as with DENconV-1 473 infections described earlier (Goic et al. 2016), result in a specific, adaptive immune response that involves the production of vcDNA by the endogenous host reverse transcriptase. We have confirmed that this vcDNA includes circular forms that can be differentially isolated from whole DNA extracts. However, for contigs that contained only unidirectional DENV-2 sequences, it was impossible to determine whether they originated from the infecting virus or from EVE. To do so, a procedure for selecting single-cell clones free of DENV-2 infection would be needed, as previously recommended (Flegel, 2020, 2022). This would allow the individual DEN2-free single-cell clones to be separately analyzed for comparison of their cvcDNA constructs. The results from this study reveal that such a process would be successful.

Our previous work using C6/36 cells was carried out to test hypotheses regarding the shrimp response to viral pathogens because no immortal crustacean cell lines existed then or even now to do such work (Kanthong et al. 2008, 2010, Sriton et al. 2009, Gangnonngiw et al. 2010, Arunrut et al. 2011). Those earlier studies revealed the ability of C6/36 cells to individually support three different shrimp viruses (WSSV, TSV, and YHV) that remained infectious for shrimp up to 5 passages but not thereafter. On the other hand, the insect cells remained immunopositive for those 3 viruses for up to 30 split passages, and they could transfer immunopositive status to the hemocytes of challenged shrimp. This suggested the possibility of using insect cells as a source of attenuated viurses for injection into shrimp to protect (i.e., vaccinate) against disease caused by cognate viruses. A first attempt to do this with a yellow head virus (YHV) failed (Gangnonngiw & Kanthong 2023). However, the hemocytes of the injected shrimp in the failed vaccination attempt were converted to immunopositive status for YHV, and there was a significant delay in their mortality. Since the “attenuated vaccine” comprised a crude extract from whole insect cells, it is possible, based on the results described herein, that vcDNA from YHV in those extracts contributed to this delay in mortality.

In addition, we have revealed here that cvcDNA prepared *in vitro* after RCA targeting an EVE of DENV-2 can reduce CPE in naive cells challenged with DENV-2. This experiment still exhibited the same level of DENV-2 viral replication as the untreated cells, with a previous report in shrimp showing that a crude cvcDNA extracted from shrimp infected with the parvovirus IHHNV could result in a significant decrease in IHHNV replication (5). Further investigation is warranted into the specific prevention of CPE by cvcDNA treatment, with the addition of quantifying DENV-2 replication. To us, the fact that pre-exposure of the naïve C6-36 cells to the crude DENV-2 cvcDNA preparation prior to DENV-2 challenge allowed them to reach accommodation quicker than the untreated cells is worthy of further investigation. It as if its may indicate the presence of some preliminary factor(s) necessary for accommodation to occur were already present. It is somehow similar to the earlier discovery that C6-36 cells already accommodating DENV-2 could more rapidly accommodate the densovirus *Aedes albopictus* densovirus (*Aal*DNV) and vice-versa (Burivong et al. 2004). It suggests the presence of a critical factor necessarily for accommodation to occur. One possibility may be host-derived reverse transcriptase (RT) or some mechanism related to it. It is a key enzyme in the viral accommodation process, and our recent work described at BioRxiv has revealed that RT inhibition with cells that have accommodated DENV-2 revert to the CPE state (Gangnonngiw et al. 2025).

These results have shown that C6/36 cells may be used as a convenient model to study the detailed mechanisms by which cvcDNA is produced directly from viral RNA and whether these mechanisms differ from those that give rise to cvcDNA from EVE. We hypothesize that C6/36 cells or other immortal insect cell lines may also produce cognate cvcDNA and EVE when challenged with shrimp viruses. If so, they might provide a convenient tool to develop EVE-clone libraries to facilitate screening for the most protective EVE or combinations thereof for use with shrimp. This is already possible, of course, for insect viruses, in general, using their matching immortal cell lines.

## ACKNOWLEDGEMENTS

This research project was supported by Rajamangala University of Technology Tawan-ok ( Basic Research Fund: fiscal year 2021, **Project no. 50063**) and Mahidol University ( Fundamental Fund: fiscal year 2023 by National Science Research and Innovation Fund (NSRF) **Grant no. FF-056/2566**).

## Supplementary information

**Yellow highlights indicate Dengue virus type 2 complete genome sequences (M29095.1).**

**Turquoise highlights indicate Aedes albopictus isolate C6/36 genome fragments.**

**Green highlights indicate repeat regions representing circular DNA.**

>k141_157363 flag=0 multi=3.5761 len=417

BLASTN output showed that the positions of k141_157363, which matched the Dengue virus type 2 complete genome (M29095.1), were 1-147 (147 bp) and 147-281 (135 bp).

BLASTN output showed that the position of k141_157363, which matched the Aedes albopictus isolate C6/36 (NW_017856292.1), was 278-417 (140 bp).

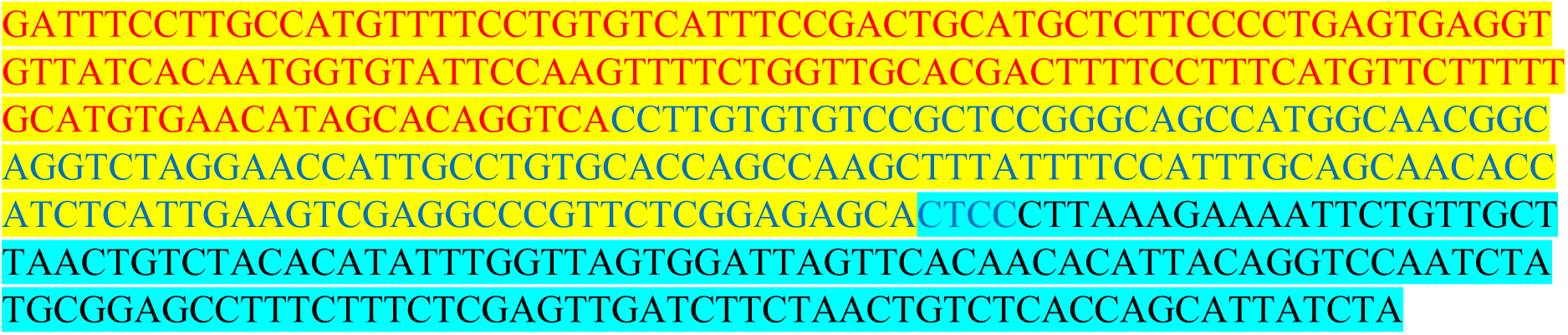

>k141_42362 flag=1 multi=1.0000 len=298

BLASTN output showed that the position of k141_42362, which matched the Dengue virus type 2 complete genome (M29095.1), was 159-298 (140 bp).

BLASTN output showed that the position of k141_42362, which matched Aedes albopictus isolate C6/36 (NW_017858069.1), was 1-140 (140 bp).

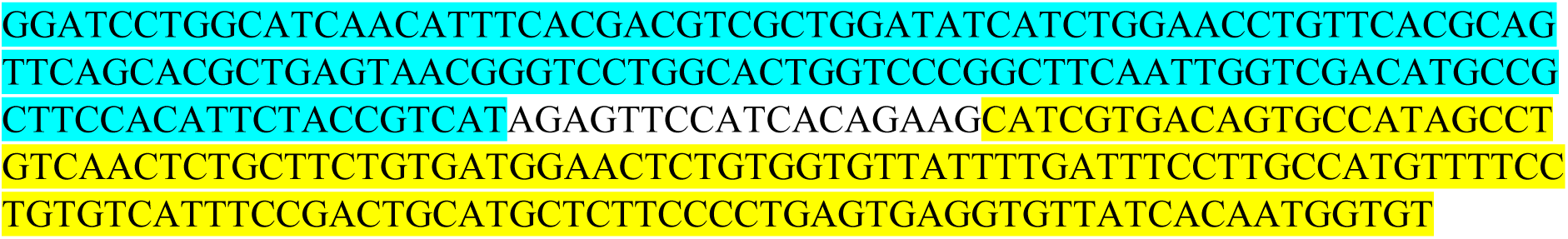

>k141_43783 flag=0 multi=7.4781 len=2061

BLASTN output showed that the positions of k141_43783, which matched the Dengue virus type 2 complete genome (M29095.1), were 1-141 (141 bp), 141-387 (247 bp), and 1179-2061 (884 bp).

BLASTN output showed that the positions of k141_43783, which matched the Aedes albopictus isolate C6/36 (NW_017856292.1), were 387-1031 (645 bp), 1028-1122 (95 bp), and 1110-1194 (85 bp).

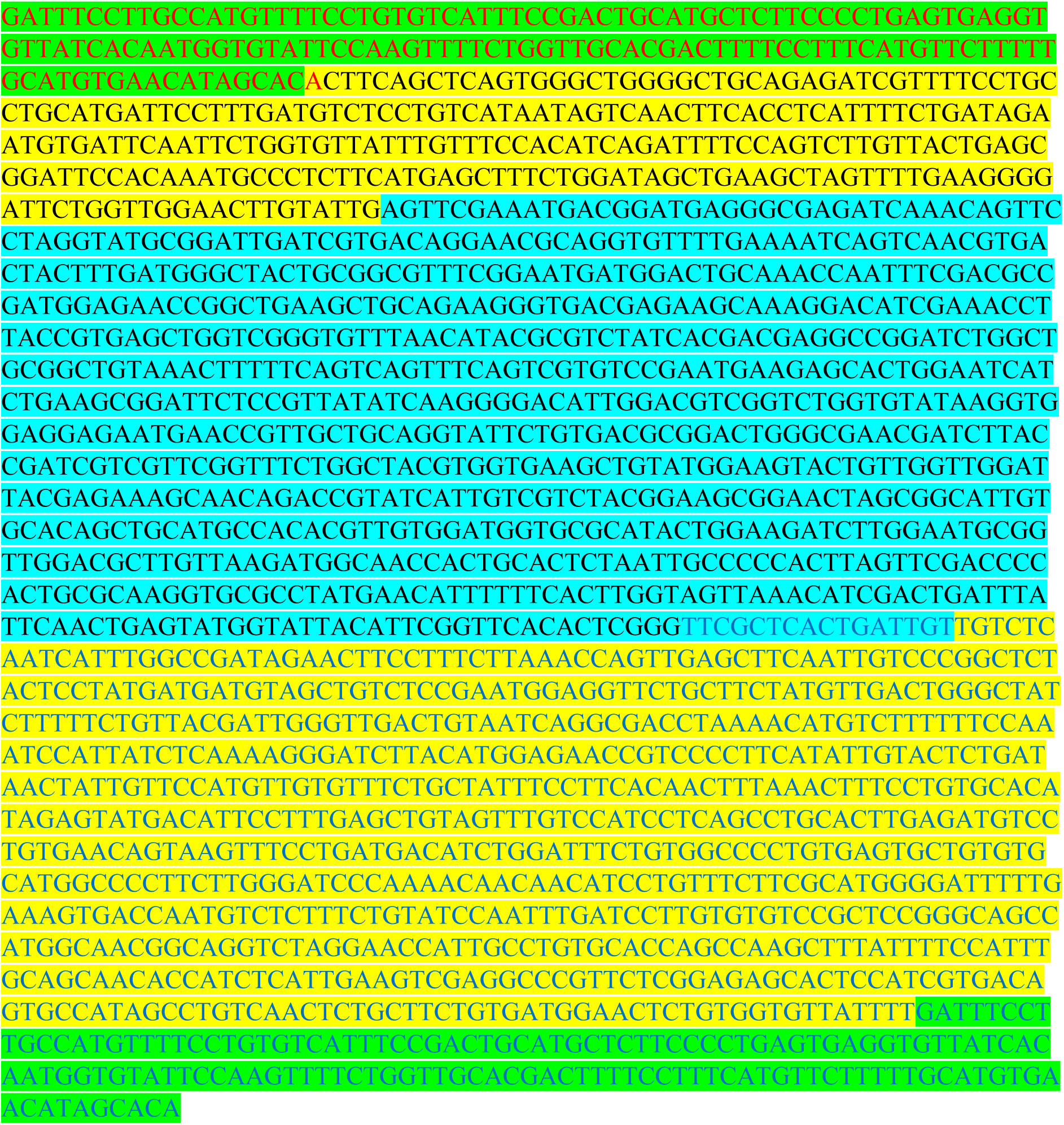

>k141_554965 flag=3 multi=22.0615 len=499

BLASTN output showed that the positions of k141_554965, which matched the Dengue virus type 2 complete genome (M29095.1), were 4-96 (93 bp), 95-272 (178 bp), 277-361 (85 bp), and 362-454 (93 bp).

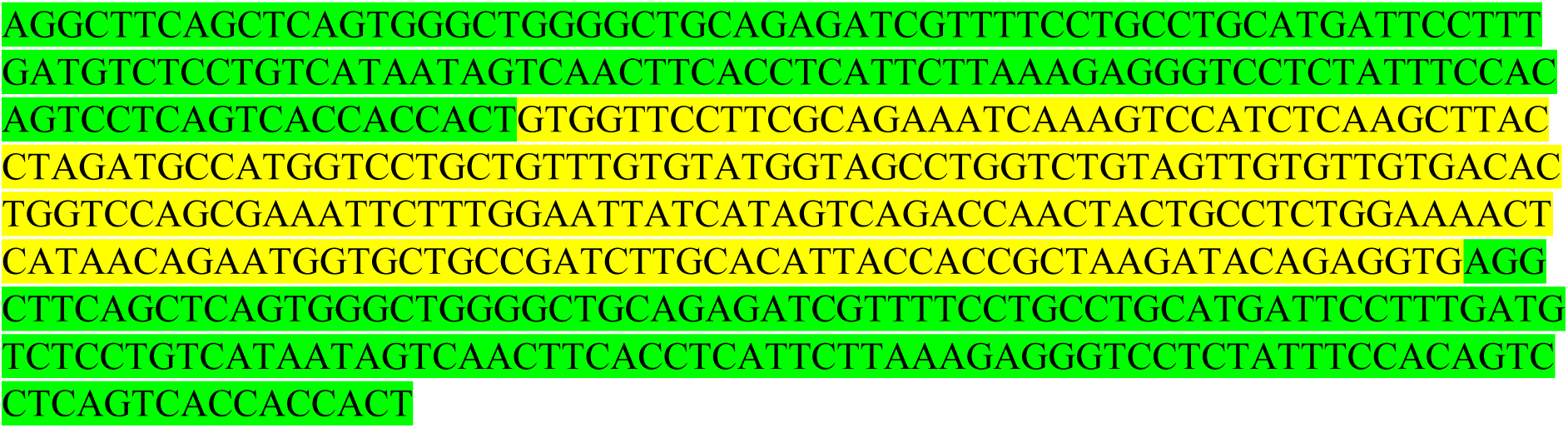

---

**Supplementary Figure S1.**
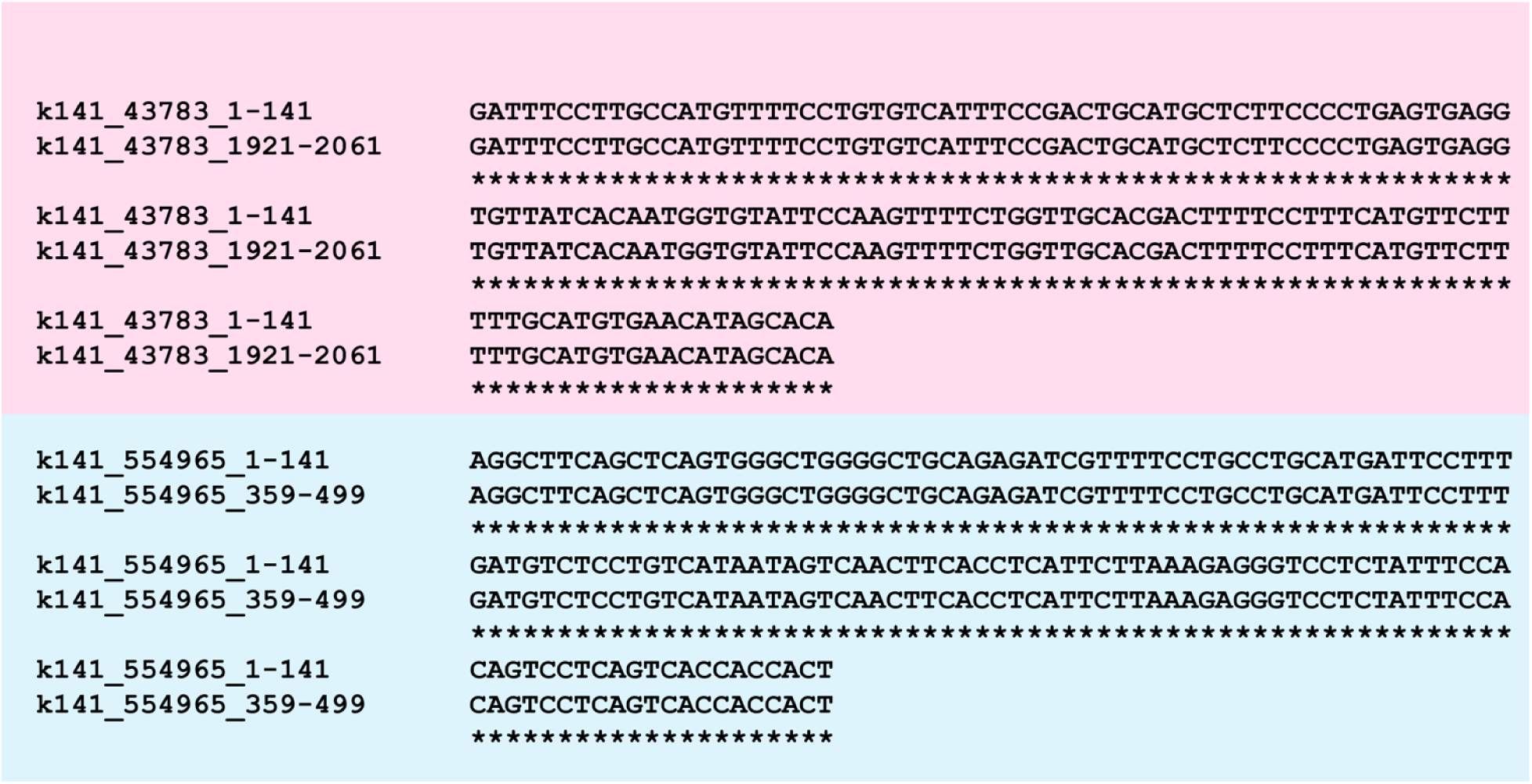
141 bp overlap regions using sequence ailgnment

**Other supplementary information available upon request.**

